# A rapid and reliable multiplexed LC-MS/MS method for simultaneous analysis of six monoamines from brain tissues

**DOI:** 10.1101/2021.05.24.445477

**Authors:** Sejal Davla, Edward Daly, Jenn Nedow, Ari Gritsas, Laura Curran, Lorne Taylor, Donald J. van Meyel

## Abstract

Monoamines are a class of neuromodulators that are crucial for a variety of brain functions, including control of mood, movement, sleep and cognition. From mammals to insects, the nervous system is enriched in monoamines such as dopamine, serotonin and melatonin, analytes which range from being highly polar to non-polar. Here we developed a method using liquid chromatography coupled with mass spectrometry (LC-MS) to quantify in a single run the amounts of six distinct monoamines in extracts from dissected *Drosophila* and mouse brain tissues. The measured monoamines were dopamine (DA), serotonin (also known as 5-hydroxytryptamine (5-HT)), octopamine (OA, an insect equivalent of norepinephrine), tyramine (TA), melatonin (MT) and N-acetyl-hydroxy-serotonin (NAS). The analytical range of these monoamines was between 0.25 to 5.0 ng/mL.

## Introduction

Biogenic monoamines are low molecular weight bases with a range of chemical polarities. Monoamines are involved in numerous biological functions in microbes, plants and animals. Found in the nervous systems of all animals, including insects, rodents and humans, certain monoamines are neuromodulators that regulate neural circuits for various physiological processes such as feeding, locomotion, sexual behavior, sleep/arousal and learning and memory (1). There has been considerable interest in the quantitative analysis of some of the major monoamines such as dopamine and serotonin, as changes in their levels are thought to contribute directly to a variety of neurological pathologies, including Parkinson’s disease, depression and sleep disorders (2, 3). A popular method to measure monoamines in tissues involves high-performance liquid chromatography coupled with electrochemical detection (HPLC-EC) (4-6). However, this technique is limited because other charged compounds in tissues can interfere with monoamine detection, and because the high oxidation potential required for this method restricts sensitivity for monoamines that are present in small quantities (7). To address these limitations, mass spectrometry (MS) has become a preferred mode of detection because of its high degree of selectivity, specificity and sensitivity. MS assays for monoamines (also known as biogenic amines) have previously incorporated a negative ion gas chromatographic (GC-MS) mode (8, 9) or, more commonly, a positive ion liquid chromatography tandem mass spectrometry (LC-MS/MS) mode (10). Several studies have measured biogenic amines from a range of biological samples that do not require exhaustive extraction, such as cerebrospinal fluid, plasma, serum, urine and saliva (11-13). Some studies have used different extraction protocols and solvent combinations to measure monoamines and their metabolites, either serially or simultaneously, from the rodent brain or intact nematodes (14-16). However, each of these methods has been limited either because they focus only on the dopamine and serotonin pathways, or because they are semi-quantitative of only the relative abundance of monoamines in a sample. In addition, high-resolution instrumentation capable of accurate mass assignment has been used to assess biogenic amine levels within single cells from the brains of fruit flies (*Drosophila* melanogaster), an important and well-studied genetic model system (17). Most applications in biomedical research involve homogenized tissues to measure both extracellular and intracellular levels of monoamines, and they are often performed in parallel runs that can exhaust precious biological specimens. Here, we describe a new multiplexed LC-MS/MS method that incorporates the use of a Triple Quadrupole Mass Spectrometer (QQQ) for the simultaneous, accurate, and specific measurement of dopamine (DA), serotonin (also known as 5-hydroxytryptamine (5-HT)), octopamine (OA, an insect equivalent of norepinephrine), tyramine (TA), melatonin (MT) and N-acetyl-hydroxy-serotonin (NAS), an intermediate in the synthesis of MT from 5-HT (18).

## Materials and Methods

### Chemicals

The reference standards for DA, 5-HT, OA, TA, MT, and NAS were purchased from Sigma (Mississauga, Canada or St. Louis, MO, USA). The deuterated standards d4-DA, d4-5-HT, d3-OA, d4-TA, d4-MT and d3-NAS were purchased from CDN Isotopes, Montreal, Canada. LC/MS grade solvents acetonitrile (ACN), ethanol (EtOH) and formic acid (FA) were purchased from Fisher Scientific. Type 1 water (18 mΩ resistance) was produced on site as needed.

### Lysate preparation from brain tissues

For *Drosophila*, each sample was comprised of 20 brains from one-to two-week old adults that were dissected in ice-cold 1X phosphate-buffered saline (PBS) solution, then centrifuged and the PBS removed. For mouse, each sample was comprised of 0.1 mg of dissected brain cortex tissue (C57BL/6J strain). Mice were euthanized by intraperitoneal injection of ketamine anesthetic cocktail (ketamine: 100mg/kg, xylazine: 10mg/kg, acepromazine: 3mg/kg) and then perfused with 1X Dulbecco’s Phosphate Buffer Saline (Gibco). All mouse procedures were carried out according to Animal Use Protocol # 2001-3995 approved by McGill University and its Affiliated Hospitals’ Research Institutes, following standards established by the Canadian Council on Animal Care. Mice used in this study were euthanized for experiments collecting their sciatic nerves, and the brain tissues used here would have been discarded otherwise. Fly or mouse brain tissues were immediately homogenized in 50µL (fly brains) or 200µL/500µL (mouse brain) of 0.1% FA solution with a motorized pestle. After centrifugation, the supernatant was collected and stored at -80°C until analysis by LC-MS/MS.

### Stock solutions

Primary stock solutions of unlabeled or deuterated monoamines were prepared in anhydrous ethanol to provide a nominal concentration of 1.0 mg/mL. The stock solutions were stored in the dark at -80°C for up to 6 months. Individual sub-stocks were prepared at 100 µg/mL in anhydrous ethanol. Further, dilutions of these sub-stocks (1µg/mL, in EtOH), were used to tune the mass spectrometer. These sub-stock solutions were stored at -20±4°C, protected from light. Two distinct working solutions were then prepared in anhydrous EtOH. The first contained all the non-labeled analytes (DA, 5-HT, OA, TA, MT and NAS) at a nominal concentration of 10 µg/mL each. The second contained all the deuterated (dx) internal reference standards (IS) (d4-DA, d4-5-HT, d3-OA, d4-TA, d4-MT and d3-NAS). Both of these combined working solutions were then further diluted to provide a nominal concentration of 1.0 µg/mL spiking solution. To optimize chromatographic conditions, a reference solution was prepared from these two diluted (1.0 µg/mL) spiking solutions (using 10/90 EtOH/water containing 0.1% FA) to achieve final concentrations of 10.0 ng/mL for each unlabeled analyte, and 2.5 ng/mL for each deuterated IS. This reference solution was used to monitor system performance and was made fresh for each analytical run.

### LC-MS/MS

Instrumentation consisted of a Thermo-Scientific Quantiva™ triple quadrupole mass spectrometer (QQQ) which incorporated a heated electrospray ionization (HESI) source, a Thermo-Scientific UltiMate™ 3000 UHPLC system (including UltiMate™ 3000 RS autosampler). The HESI source voltage was 3200 V, the Sheath gas 30 L/min, Aux gas 10 L/min, Sweep gas 2 L/min, the Ion Transfer Tube temperature was 325°C, the Vaporizer temperature was 350 °C, the CID gas was 1.5 (mTorr) and the dwell time was 50 msec. The Q1 resolution full width half maximum (FWHM) was 0.4 and the Q3 Resolution (FWHM) was 0.7. The instrument ran in a selected reaction monitoring (SRM) positive ion detection mode at various collision energies (CE in V) for the mass transitions outlined in Table 1. The MS was auto-tuned as per instrument manufacturer’s specifications.

**Table 1:** Optimized Mass transitions and Corresponding Collision Energies.

| Analyte | Mass<br>(M+H<sup>+</sup>) | Q1/Q3 masses | Collision<br>energy<br>(volts) |
| --- | --- | --- | --- |
| DA | 154 | 137.1/91.1<sup>1</sup> | 17.1 |
| 5-HT | 177 | 160.1/115.1<sup>1</sup> | 22.7 |
| OA | 154 | 136.1/91.1<sup>2</sup> | 19.1 |
| TA | 138 | 138.1/121.0 | 10.2 |
| MT | 233 | 233.1/174.1 | 14.0 |
| NAS | 219 | 219.1/160.1 | 14.5 |
| d4 -DA | 158 | 141.1/95.1<sup>1</sup> | 18.5 |
| d4-5-HT | 181 | 163.9/118.1<sup>1</sup> | 25.3 |
| d3- OA | 157 | 136.3/93.1<sup>2</sup> | 18.2 |
| d4- TA | 142 | 142.1/125.1 | 10.2 |
| d4-MT | 237 | 237.1/177.9 | 13.8 |
| d3-NAS | 222 | 222.1/160.1 | 14.4 |

<sup>1</sup>A Precursor Ion of highest abundance representing a net loss of -17 u ([M-17])<sup>+</sup> from the protonated molecular ([M+H])<sup>+</sup> was chosen for the MRM transition.
<sup>2</sup>A Precursor Ion of high abundance representing a net loss of -18 u ([M-18])<sup>+</sup> from the protonated molecular ([M+H])<sup>+</sup> was chosen for the MRM transition.

Chromatography of monoamines on an Agilent Eclipse Plus™ C-18 analytical column (50mm X 2.1mm ID x 1.8 µm particle) involved adaptation of a method (19) using gradient elution of a binary solvent system that incorporated (A) 0.1% FA (aqueous) and (B) ACN + 0.1% FA as follows: 0 - 0.5 min, 5% B; 0.51 - 6.0 min, 80 % B; 6.01 - 8.0 min, 80 % B; 8.1 min - 10.2 min, 5 % B. A divert valve was used to divert the first 0.25 min to waste. The solvent flow rate was 200 µL/min. A 10µL injection volume was used. The MS/MS acquisition time was 12.1 min - the total run time was 14.1 min (allowing a 2 min column reconditioning step).

The chromatographic system was determined to be stable and reproducible by injecting a reference solution, at a nominal concentration of 5.0 ng/mL (50 pg on column, each analyte), three times at the beginning and at the end of the chromatographic run. The percent difference of the mean integrated peak areas injected reference standards was ≤ 25% CV. No carry-over was observed, as assessed by injecting a blank sample following the injection of the reference solution.

### Calibration Curve Preparation

Solutions outlined in Table 2 were used to spike 1.0 mL buffer (10/90 EtOH/water containing 0.1% FA) at final concentrations of 5.0, 2.5, 1.0, 0.5, 0.25 and 0 ng/ml for the calibration curve (n=3). The Lower Limit of Quantification (LLOQ) of the analytical curve was based on the detector response of NAS (0.25 ng/mL) which had the lowest quantifiable chromatographic peak area response (3:1, S:N) relative to the other analytes being surveyed. A double blank solution control (no analyte, and no dx-IS), and a Standard A (dx-IS only) were also prepared and included with the calibration curve. Results used for the standard curve are shown in Table 3.

**Table 2:** Concentrations of spiked standards in solution calibration curve analyzed by Thermo Quantiva LC-MS/MS (QQQ)

| Standards | Volume of combined analyte spiking solution used (μl)<sup>1</sup> | Volume of combined internal standard analyte spiking solution used (μl)<sup>2</sup> | Final concentration of combined analytes (ng/ml)<sup>3</sup> | Final concentration of combined internal standard (ng/ml)<sup>3</sup> |
| --- | --- | --- | --- | --- |
| Double Blank | 0.0 | 0.0 | 0.0 | 0.0 |
| Standard A (IS only) | 0.0 | 25 | 0.0 | 2.5 |
| Standard B<br>(LLOQ) | 25* | 25 | 0.25 | 2.5 |
| Standard C | 50* | 25 | 0.5 | 2.5 |
| Standard D | 10** | 25 | 1.0 | 2.5 |
| Standard E | 25** | 25 | 2.5 | 2.5 |
| Standard F | 50** | 25 | 5.0 | 2.5 |

- (1) Combined analyte spiking solution concentrations: (\*) = 1000ng/ml, (\*\*) = 100ng/ml, in EtOH, spiked into 1.0 mL final buffer (10/90 EtOH/water plus 0.1% formic acid). Total volume for each standard = 1 mL. - (2) Combined deuterated IS spiking solution concentration: 1000ng/ml in EtOH, spiked into 1.0 mL final, buffer (10/90 EtOH/water plus 0.1% formic acid). Total volume for each standard - 1ml. - (3) Concentration of individual analytes or individual dx-IS in buffer at the various levels of the Calibration Curve.

**Table 3:**
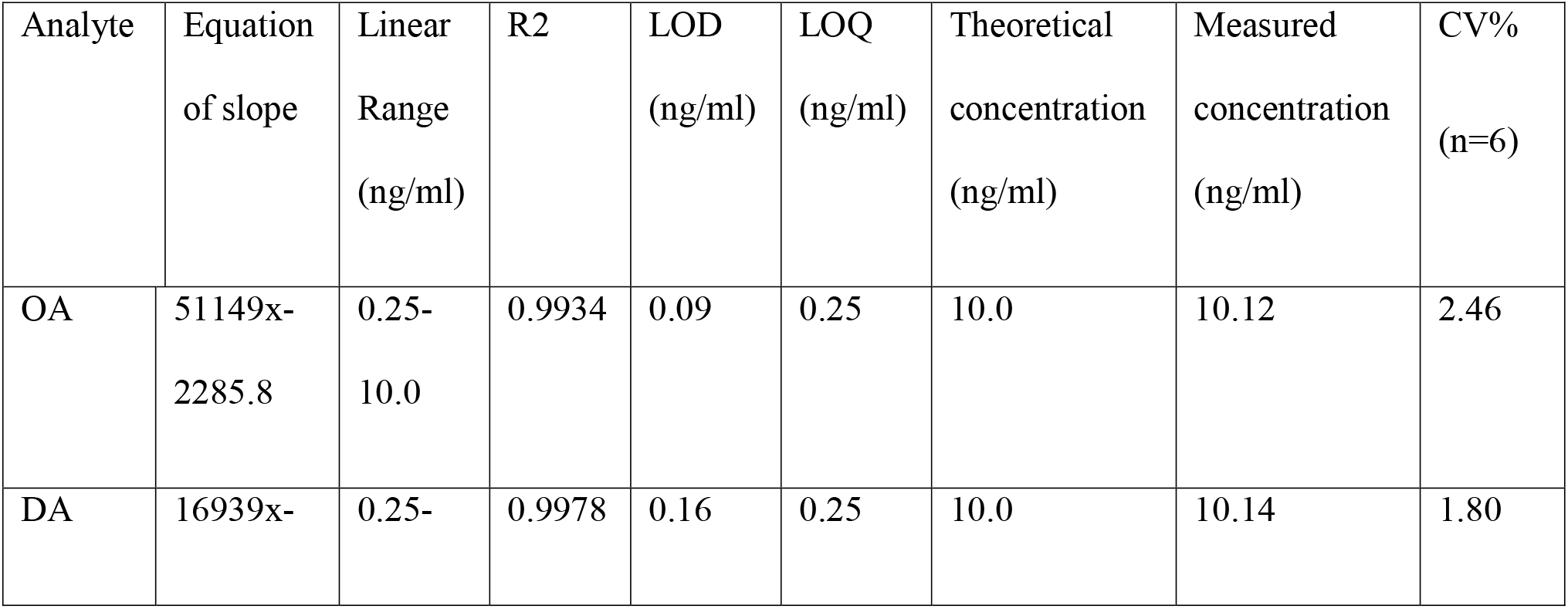
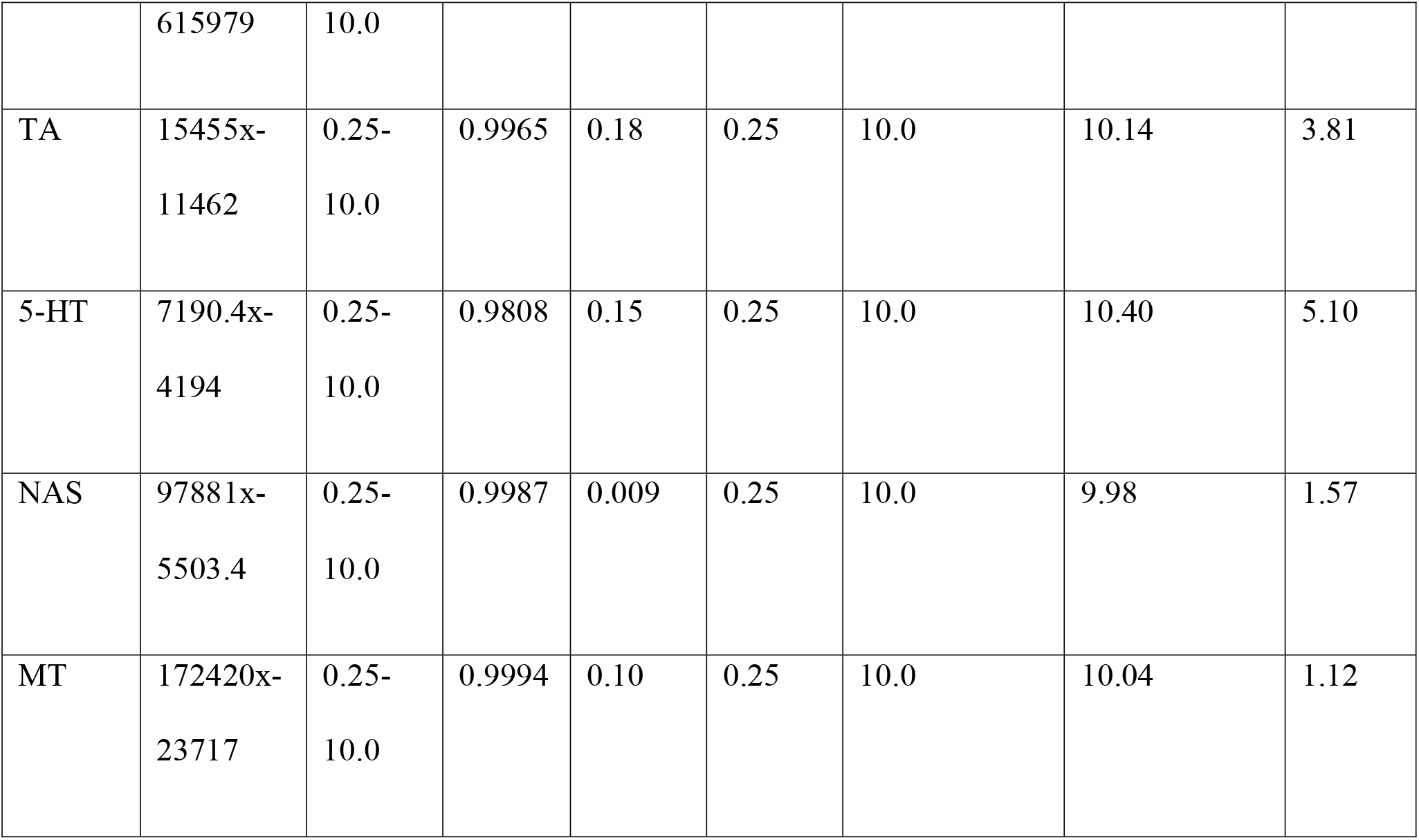
Results for linear range calibration curve.

### Sample Extraction

A 10.0 µL aliquot from each extract was introduced into the LC-MS/MS system. Extraction recovery was determined by spiking an extract for a nominal concentration of 1.0 ng/mL with the diluted (1.0 µg/mL) combined spiking solution and comparing with the equivalent concentration spiked in solution. The apparent extraction recovery was greater than or equal to 85% for the individual analytes.

## Results and Discussion

Monoamines are important neuromodulators, and it is desirable to measure several in a single analytical run to save time and brain tissue from precious specimens. Monoamines are grouped based on their chemical structure, such as indoleamines (e.g. 5-HT, MT, NAS), catecholamines (e.g. DA), and phenethylamines (e.g. OA, TA). Their range of polarities presents a challenge for their separation from one another in a single sample, and so we developed a LC-MS/MS method that incorporates gradient reverse-phase chromatographic elution of extracts from dissected brains of flies and mice, two commonly used model systems for biomedical research. We coupled that with mass spectrometric analysis of deuterated standards to accurately quantify these six monoamines at once.

### Analysis of synthetic reference standards

Analytical conditions were originally developed using a Reference Standard in solution containing DA, 5-HT, OA, TA, MT and NAS which included the corresponding deuterated internal standards - d4-DA, d4-5-HT, d3-OA, d4-TA, d4-MT and d3-NAS. The concentration of the unlabeled standards was originally 10 ng/mL for each analyte (100 pg on column), and subsequently lowered to 5.0 ng/mL and 2.5 ng/mL for the labeled standards. The precursor ions chosen for the MRM mass transitions representing OA and DA were ([M-17]^+^) ions, which avoid isobaric masses representing the ([M+H]^+^) of either analyte.

We achieved adequate chromatographic separation of the six monoamines in a single analytical run by incorporating 0.1% FA (Aq) in mobile phase A. We applied it to all subsequent fly and mouse brain extracts. For standards in solution, we produced representative SRM reconstructed total-ion current chromatograms (RTICC, Fig 1) and report optimized mass transitions and collision energies for DA (mass-to-charge ratio transition (m/z) 137→91), 5-HT (m/z 160→115), OA (m/z 136→91), TA (m/z 138→121), MT (m/z 233→174) and NAS (m/z 219→160) (Table 1). These analytes eluted from the column as unique individual chromatographic entities: OA eluted first with a Retention Time (RT) of 0.60 min, then DA (RT 0.72 min), TA (RT 0.83 min), 5-HT (RT 1.20 min), NAS (RT 3.68 min) and MT (RT 4.62 min) (Fig 1). Chromatographic resolution was not obtained for enantiomers of DA, OA and TA. For 5-HT, a chromatographic peak eluted at RT 3.67 min, corresponding to the gas phase loss of the amino-acetyl group ([M-59]^+^) from NAS, resulted in ions detected at m/z 160. SRM mass chromatograms for corresponding six deuterated monoamines in the internal standard solution were produced for comparison (Fig 2).

**Figure 1.**
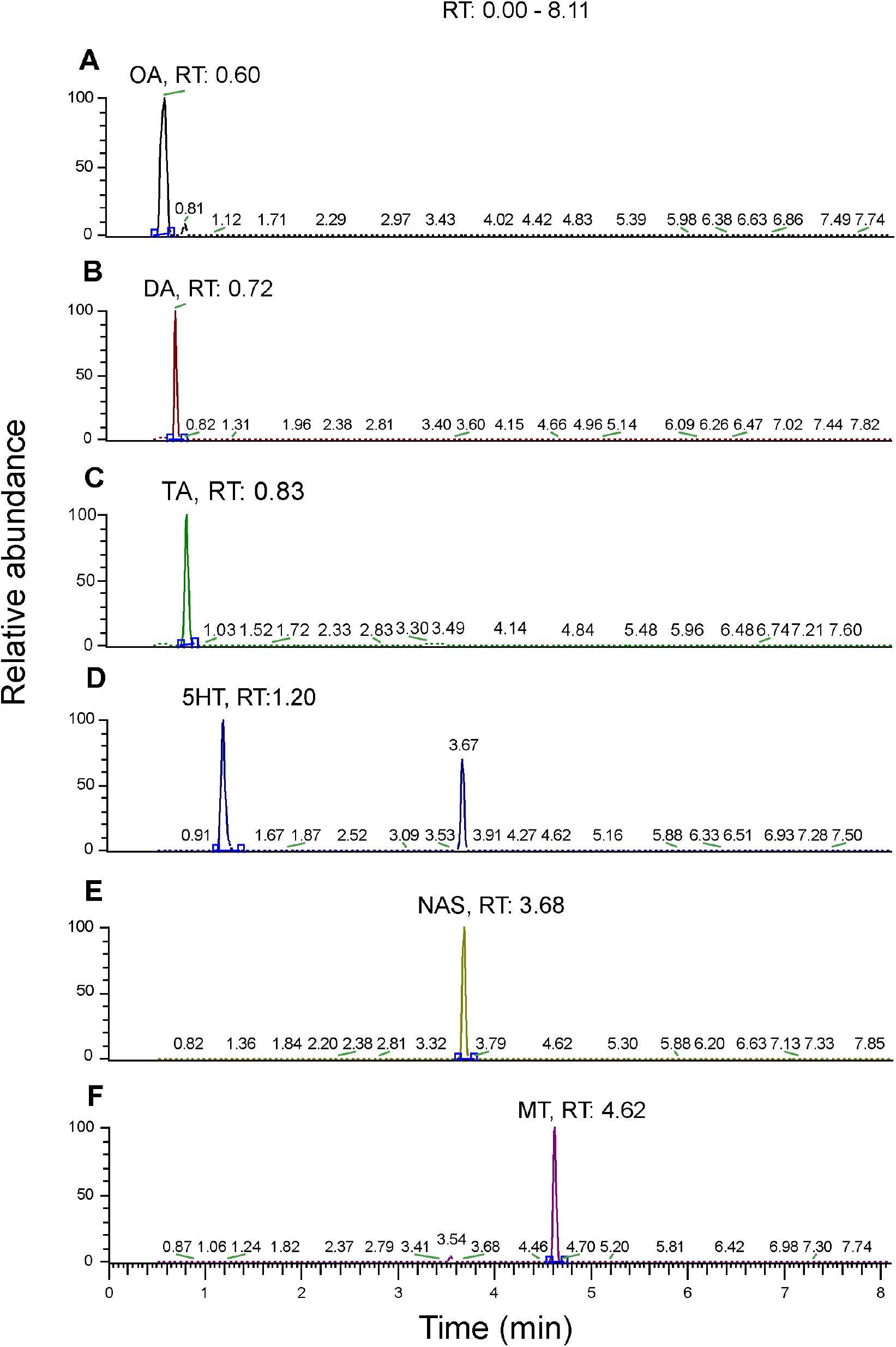
Representative traces for mass spectrometric analysis of reference standards. SRM scan HESI-MS/MS mass chromatograms of a reference standard (10 ng/mL, 1000 pg ‘on column’) in solution, 10uL injection; (A) OA, (m/z 136→91), (B) DA (m/z 137→91), (C) TA (m/z 138→121), (D) 5-HT (m/z 160→115), (E) NAS (m/z 219→160) and (F) MT, (m/z 233→174).

**Figure 2.**
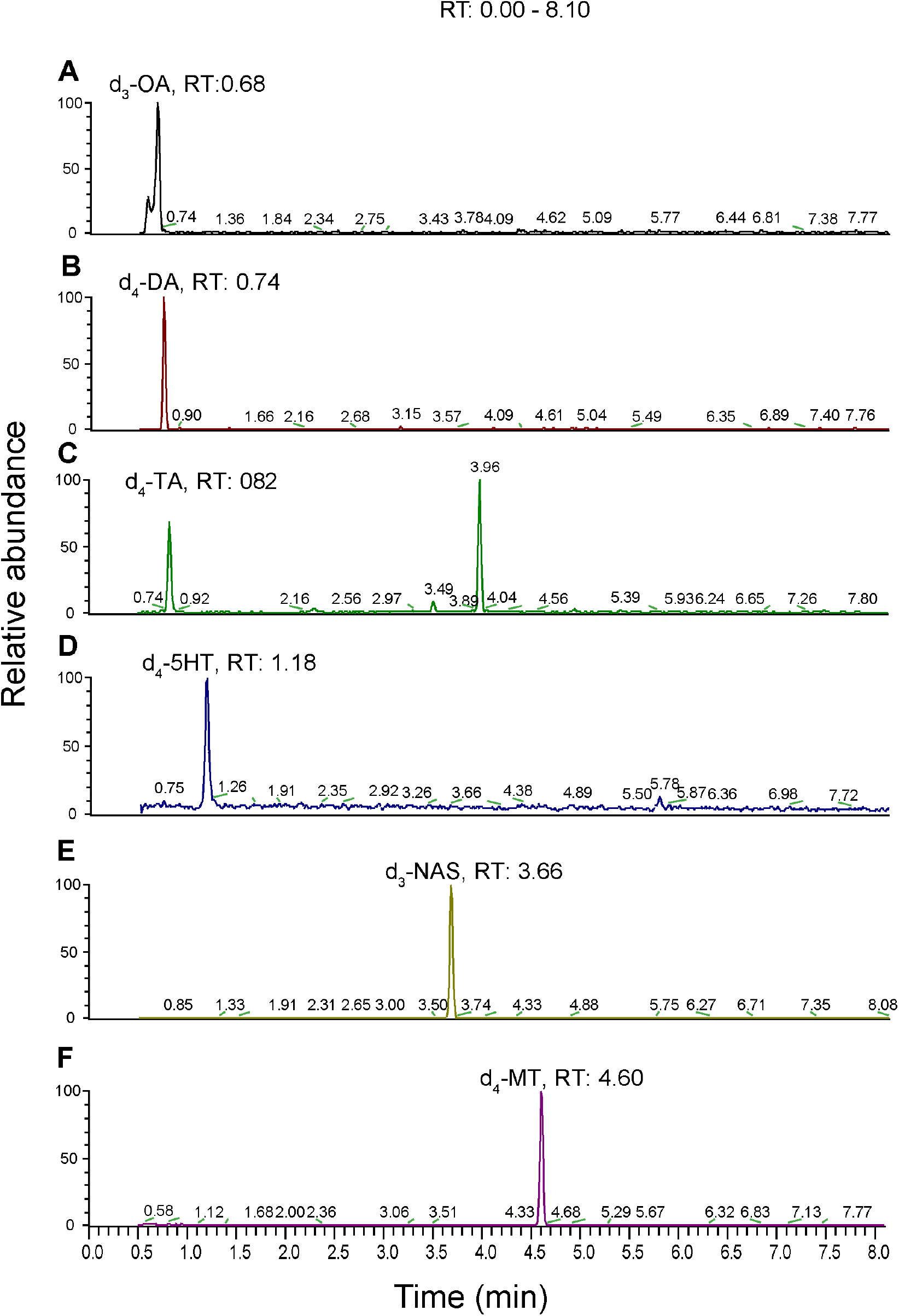
Representative traces for mass spectrometric analysis of deuterated reference standards. SRM scan HESI-MS/MS mass chromatograms of a reference standard (10 ng/mL, 1000 pg ‘on column’) in solution, 10uL injection; (A) d3-OA, (m/z 139→93), (B) d4-DA (m/z 141→95), (C) d4-TA (m/z 141→125), (D) d4-5=HT (m/z 164→118), (E) d3-NAS, (m/z 222→160) and (F) d4-MT, (m/z 237→178).

### Detection of monoamines in extracts from *Drosophila* brains

We then tested extracts of dissected adult *Drosophila* brains and produced representative SRM mass chromatograms (Fig 3). Under these conditions trace levels of OA were detected at RT 0.58 min, then moderate levels of DA (RT 0.78 min), TA (RT 0.85 min), 5-HT (RT 1.22 min), and finally trace levels of NAS (RT 3.68 min). As noted above, a peak eluting at RT 3.67 min was found in mass channel associated with 5-HT, likely corresponding to the gas phase loss of the amino-acetyl group ([M-59]^+^) group from NAS. This potential fragmentation of NAS might account for the low abundance of NAS detected in the fly head extracts. MT was expected to elute at RT 4.73 min and was detectable in trace amounts in the fly brain lysates we tested.

**Figure 3.**
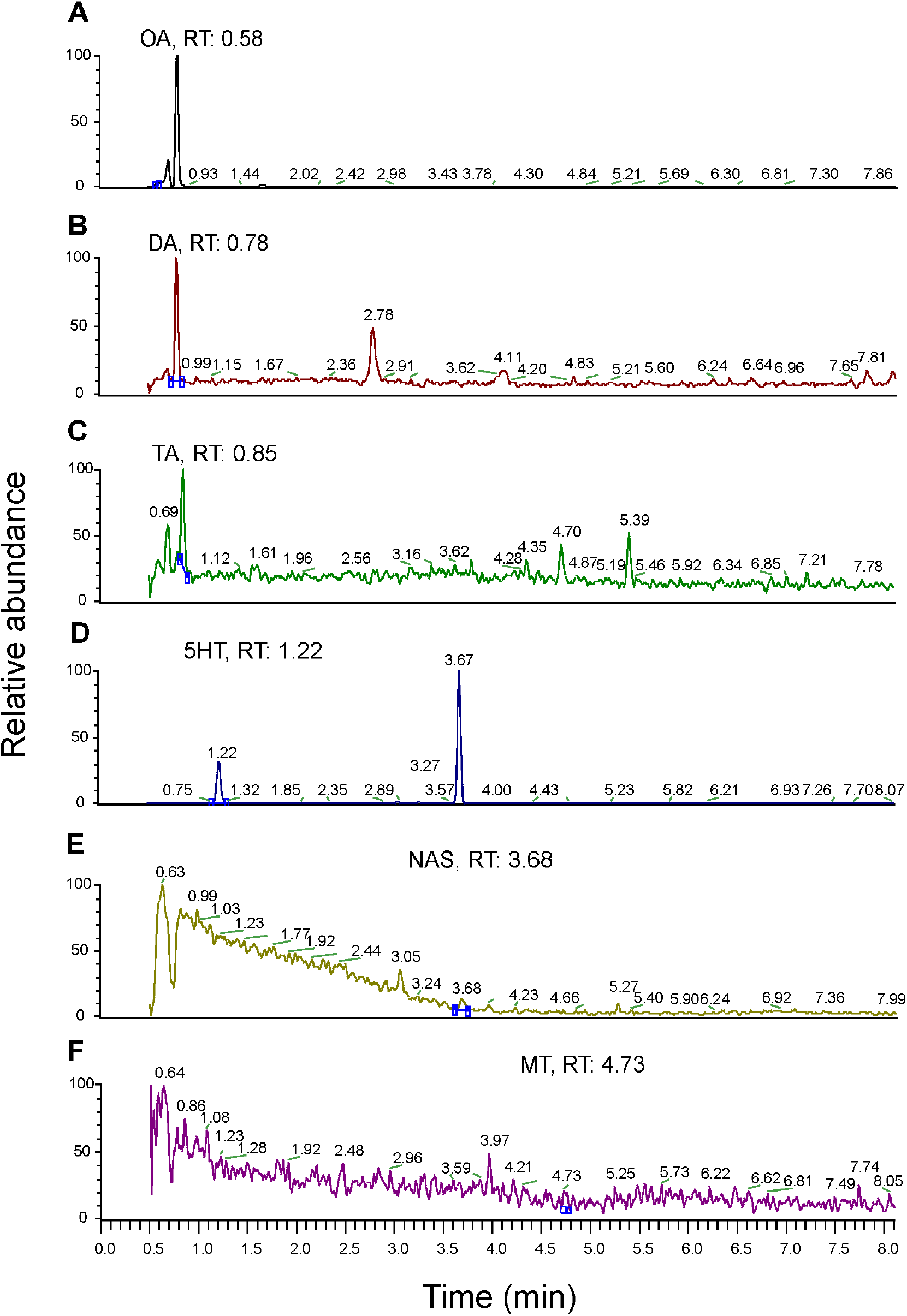
Representative traces for mass spectrometric analysis of monoamines derived from brain lysates. SRM Scan HESI-MS/MS Mass Chromatograms of Fly head extract A2 10uL injection; (A) OA, (m/z 136→91), (B) DA, (m/z 137→91), (C) TA (m/z 138→121), (D) 5-HT (m/z 160→115), (E) NAS (m/z 219→160) and (F) MT, (m/z 233→174).

### Quantification of monoamines in extracts from *Drosophila* brains

To obtain standard curves for quantification, samples of 1 ml of each analyte DA, 5-HT, OA, TA, NAS, and MT were prepared in the concentration range of 0.25 ng/mL to 5.0 ng/mL (Table 2). For analysis, each concentration was measured with three replicates and the average ion intensity ratios were calculated. The experimentally determined LLOQ was 0.25 ng/ml for each analyte (Fig 4).

**Figure 4.**
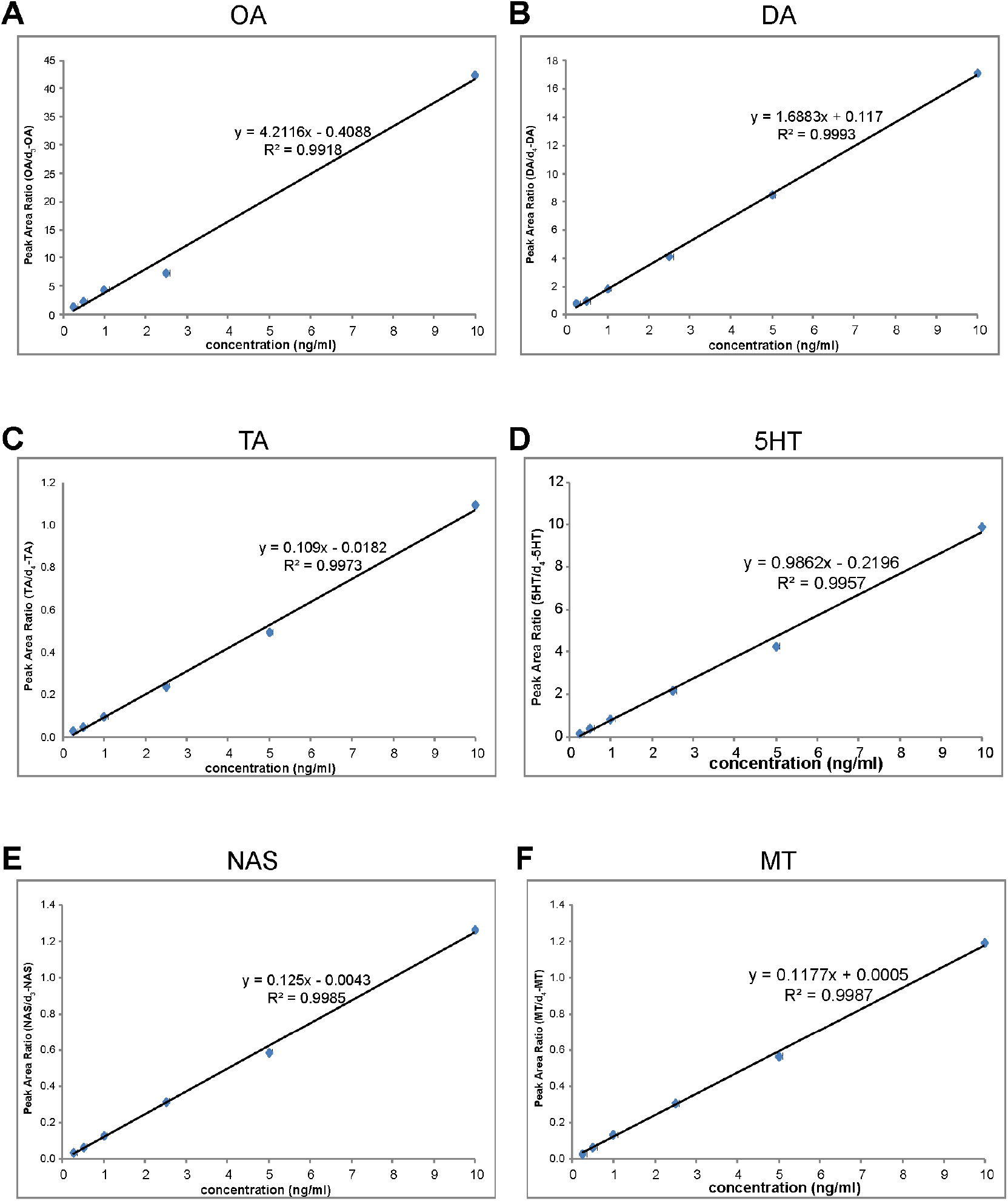
Calibration curves for quantification in MS mode of monoamines in solution: (A) OA, (B) DA, (C) TA, (D) 5-HT, (E) NAS and (F) MT. For each monoamine, peak area ratios (of unlabeled standard/d-IS) are expressed as a function of increasing concentration of the unlabeled standard. For OA, the result for 5.0ng was removed for technical reasons.

Brains of adult flies were collected in a 3-hour window each morning to control for potential circadian variability in brain monoamine levels. Each analyte eluted from the column with a RT corresponding to the RT determined for the analytical standards and associated deuterated IS. We found DA, 5-HT, TA, and NAS at moderate quantities (pg/mg brain tissue) (Fig 5). OA was detected in trace amounts: the integrated peak areas were difficult to measure against a deuterated internal standard (Fig 5). As a result of this, the relative rather than absolute levels are reported for OA.

**Figure 5.**
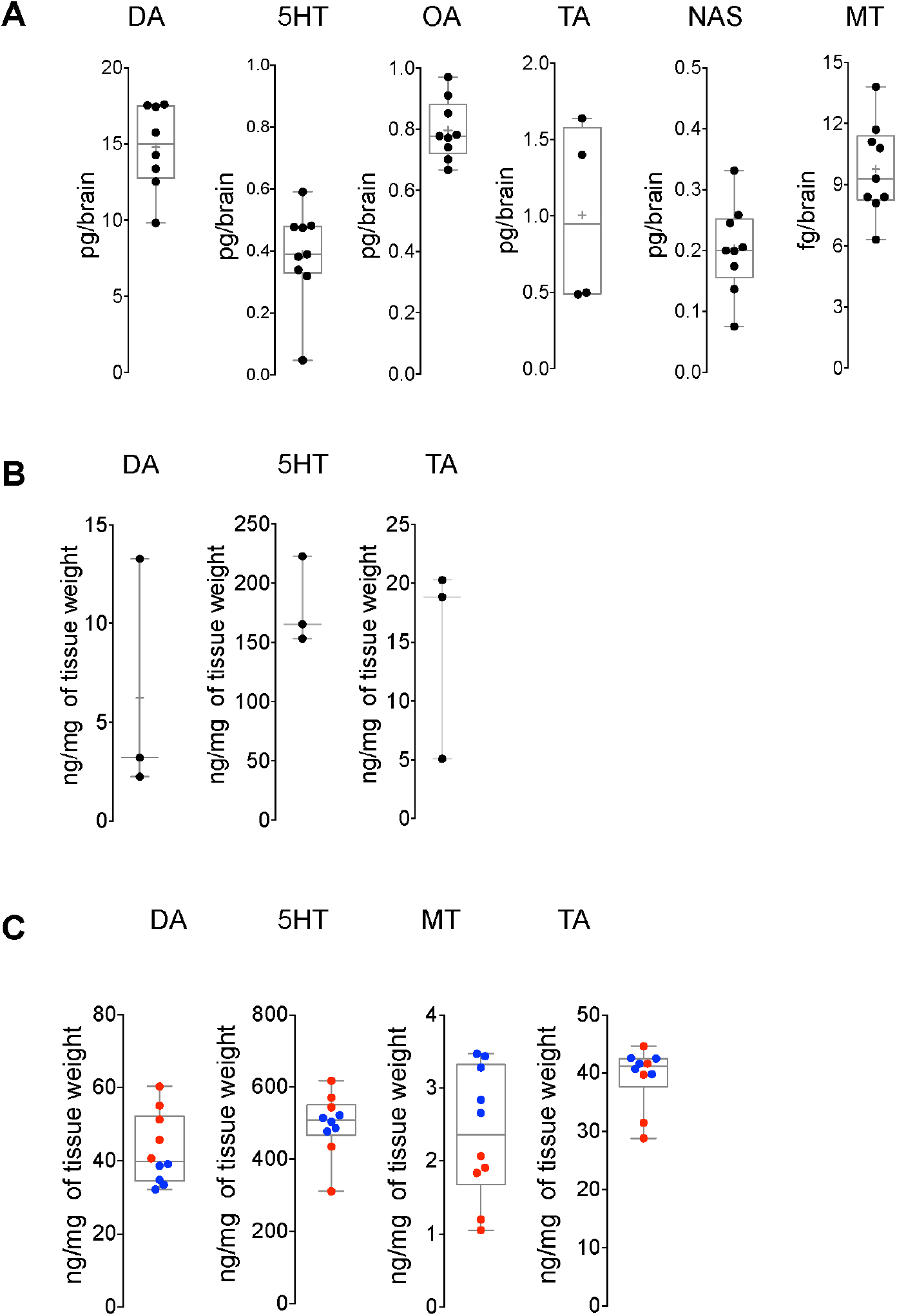
Quantitative measurements of monoamines from the fly and mouse brain lysates, where all box and whisker plots show 25-75% interquartile range (box), minimum and maximum (whiskers), median (horizontal line in box), and mean (+). (A) HPLC-MS measurement of DA (n=8), 5-HT (n=9), OA (n=9), TA (n=4), NAS (n=9), and MT (n=9) obtained from fly brains under control conditions. (B) HPLC-MS measurement of DA (n=3), 5-HT (n=3), and TA (n=3) obtained from 200µl sample extract from cortex of mouse brain under control conditions. (C) HPLC-MS measurement of DA (n=10), 5-HT (n=10), MT (n=10), and TA (n=10) obtained from 500µl sample extract the cortex of mouse brain under control conditions. Two biological replicates were used shown in blue and red and each extraction volume from a biological replicate was distributed in five vials to run in MS for technical replicates.

### Detection and quantification of monoamines in extracts from mouse brains

We dissected cortices of adult mouse brains into one of two distinct volumes of 0.1%FA (200µl or 500µl), and measured levels of four monoamines (DA, 5-HT, TA and MT) in a single analytical run. Using 200µl of sample extract, we measured levels of DA, 5-HT and TA, a vasoactive bioamine in the mouse brain (Fig 5B). However, we failed to detect MT. By using 500µl instead, we extracted more of each of DA, 5-HT and TA, and could detect and measure MT also (Fig 5C). We did not attempt to detect OA (insect-specific) or NAS in mouse brain.

Biogenic amines have been commonly measured using either gas chromatography or liquid chromatography incorporating amperometric or coulometric electrochemical detection systems (19, 21). However, the use of gradient elution reverse phase chromatography can afford an alternate mode of detection that has a highly effective degree of control and reproducibility (22, 23). For highly polar monoamines such as DA, OA and TA, this approach is challenging because they elute from the analytical column quite close to the tailing edge of the solvent front. This can be accompanied by ion suppression effects than can distort the symmetry of the eluting peaks of interest and thereby affect the determination of an accurate integrated peak area required for quantitation. In this way, ion suppression and peak distortion affect consistency in quantifying polar monoamines. In this analytical method, the incorporation of a divert valve to redirect the HPLC ‘flow to waste’ sufficiently reduced ion suppression effects and consequently improved the detection of DA and OA, as determined by consistent retention and resolution of these analytes on the analytical column. The non-polar monoamines 5-HT, MT and NAS were sufficiently retained on the column and were not affected by ion suppression or other deleterious effects. With this method, the later retention time of these analytes was not accompanied by increased ‘band spreading’ that could lead to an over-estimation of the integrated peak areas. Therefore, for monoamines ranging from highly polar to non-polar, peaks were sufficiently symmetric to allow consistent integration of peak areas for quantitative purposes.

## Conclusion

This analytical method can, in a single run using *Drosophila* brain tissue, detect and measure six monoamines that range in their polarities and abundance (DA, 5-HT, OA, TA, NAS, and MT). The method was also used to detect DA, 5-HT, TA, and MT in mouse brain tissues. Because the method is both rapid and reliable, we expect it can be widely implemented by others, and that it can be adapted readily to other tissues, organisms, or other monoamines with a range in polarity.

## Supporting Information

**S1 Fig. Representative traces for mass spectrometric analysis of a blank solution**.

## Author Contributions

The manuscript was written through contributions of all authors to the experimental design, data analysis, and writing of the manuscript. All authors have given approval to the final version of the manuscript.

## Notes

The authors declare no competing financial interest.

## Acknowledgements

This work was supported by the Research Institute of the McGill University Health Centre, and by grants to D.J.v.M. from the Canadian Institutes of Health Research (CIHR), the Natural Sciences and Engineering Research Council of Canada, and the Canada Foundation for Innovation.

