## Supplementary figures for "A rapid and reliable multiplexed LC-MS/MS method for simultaneous analysis of six monoamines from brain tissues"

Figure S1

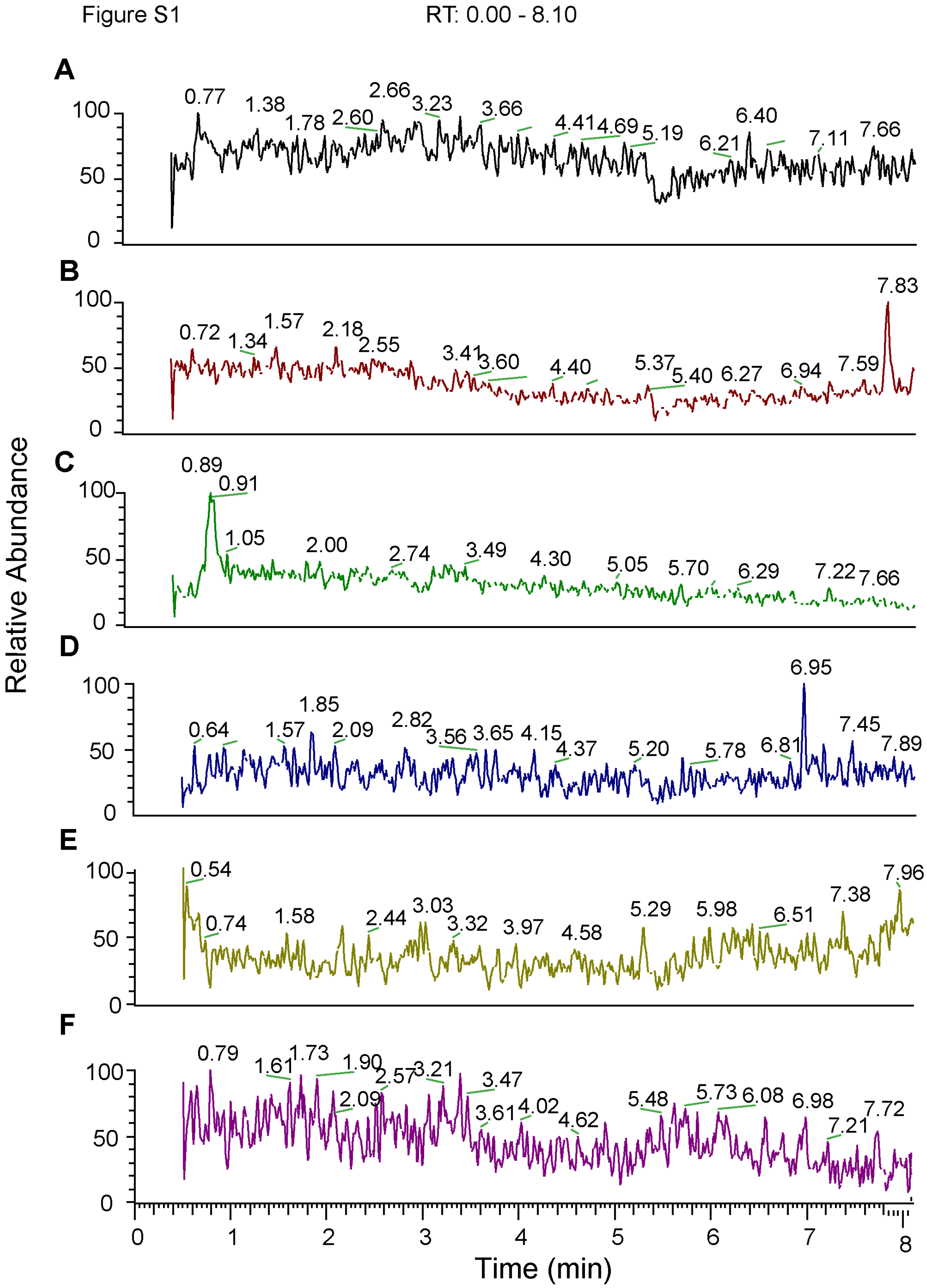


**Figure S1 Representative traces for mass spectrometric analysis of a blank solution.**

SRM Scan HESI-MS/MS Mass Chromatograms of a Blank Solution (0 ng/mL), 10uL injection;
(A) OA (*m/z* 136→ m/z 91), (B) DA (*m/z* 137→91), (C) TA (*m/z* 138→121), (D) 5-HT (*m/z* 160→115), (E) NAS (*m/z* 219→160) and (F) MT (*m/z* 233→ m/z 174).
